# High-throughput screening of more than 30,000 compounds for anthelmintics against gastrointestinal nematode parasites

**DOI:** 10.1101/2024.05.16.594481

**Authors:** Mostafa A. Elfawal, Emily Goetz, You-Mie Kim, Paulina Chen, Sergey N. Savinov, Leonard Barasa, Paul R. Thompson, Raffi V. Aroian

## Abstract

Gastrointestinal nematodes (GINs) are amongst the most common parasites of humans, livestock, and companion animals. GIN parasites infect 1-2 billion people worldwide, significantly impacting hundreds of millions of children, pregnant women, and adult workers, thereby perpetuating poverty. Two benzimidazoles with suboptimal efficacy are currently used to treat GINs in humans as part of mass drug administrations, with many instances of lower-than-expected or poor efficacy and possible resistance. Thus, new anthelmintics are urgently needed. However, screening methods for new anthelmintics using human GINs typically have low throughput. Here, using our novel screening pipeline that starts with human hookworms, we screened 30,238 unique small molecules from a wide range of compound libraries, including ones with generic diversity, repurposed drugs, natural derivatives, known mechanisms of action, as well as multiple target-focused libraries (e.g., targeting kinases, GPCRs, and neuronal proteins). We identified 55 compounds with broad-spectrum activity against adult stages of two evolutionary divergent GINs, hookworms (*Ancylostoma ceylanicum*) and whipworms (*Trichuris muris*). Based on known databases, the targets of these 55 compounds were predicted in nematode parasites. One novel scaffold from the diversity set library, F0317-0202, showed good activity (high motility inhibition) against both GINs. To better understand this novel scaffold’s structure-activity relationships (SAR), we screened 28 analogs and created SAR models highlighting chemical and functional groups required for broad-spectrum activity. These studies validate our new and efficient screening pipeline at the level of tens of thousands of compounds and provide an important set of new GIN-active compounds for developing novel and broadly-active anthelmintics.

---

Human gastrointestinal nematodes (GINs, also known as soil-transmitted helminths) include the roundworm *Ascaris lumbricoides*, the whipworm *Trichuris trichiura*, and the hookworms *Necator americanus*, *Ancylostoma duodenale*, and *Ancylostoma ceylanicum*; they are amongst the most prevalent infections of the so-called Neglected Tropical Diseases (NTDs)^1^, affecting people in the world’s poorest communities^2–4^. GINs are successful parasites of vertebrates, adapted to parasitize humans, livestock, and companion animals^5^. Morbidities associated with GIN infections in humans include anemia, malnutrition, growth stunting, cognitive impairment, pregnancy complications, and lowered immunity against other infectious diseases^2, 4, 6^. The negative impacts of GINs are hard to measure, but they are minimally responsible for > 2-3 million disability-adjusted life years (DALY)^3, 6, 7^, with hookworms accounting for almost one-half of those DALYs^3, 6^. Children who received hookworm treatment had increased school attendance and adult income^4, 8^. GINs also have enormous impacts on the health and productivity of farm animals^9^.

The World Health Organization (WHO) has recommended annual or semi-annual Mass Drug Administration (MDA) targeting preschool and school-age children to reduce worm burden and associated morbidity^10^. Four anthelmintics (levamisole, pyrantel pamoate, albendazole, and mebendazole), initially developed for livestock, are currently on the WHO list to control human GINs, with the two benzimidazoles, albendazole and mebendazole, most widely used because of their better efficacies and ease of dosing^11, 12^. In companion and livestock animals, GINs are controlled via frequent and routine treatment using a repertoire of anthelmintics, which has led to widespread and sometimes multi-drug anthelmintic resistance^13–16^. Benzimidazole resistance is well-established in veterinary parasites^17–20^. Single nucleotide polymorphisms (SNPs) in the beta-tubulin gene linked to the benzimidazole resistance in veterinary parasites have been found in human GINs^21–24^, indicating that with high drug pressure, there is likely to be an emergence for benzimidazole resistance in human GINs. In Haiti and Kenya, the frequency of benzimidazole-resistance codon 200 in *T. trichiura* was significantly increased following local MDA programs^22^. There are also multiple reports of lowered efficacy of benzimidazoles against human GINs and warnings against drug resistance^25–27^. Notwithstanding these challenges, the WHO has set an ambitious goal to eliminate human GINs by 2030^10^. Therefore, potent and broadly active anthelmintics with new modes of action are urgently needed for long-term control and elimination. One promising new drug is emodepside, which was discovered >30 years ago ^28^. However, it has not yet been approved for human use, and, as with all single anthelmintics, resistance is a concern^29, 30^. Given the lead time for new drugs to enter the clinic, it is urgent that new lead anthelmintics be put into the pipeline.

Because understanding of these parasites’ biology is still limited, whole organism phenotypic screening remains a preferred approach for anthelmintic drug discovery^31, 32^, although more recently some more directed screens, *e.g.*, based on detailed knowledge of metabolic pathways, have shown promise^33–35^. Phenotypic screening using GIN parasites for anthelmintic has included screening against *ex vivo* adult parasites, egg hatch assay, larval stages, and infective third-stage larvae^32^. However, regarding GINs that infect humans, the throughput of these screens has been limited^32, 36–38^. The free-living nematode *Caenorhabditis elegans* has been used as a surrogate model for GINs that infect humans^32, 39–42^. However, we found limitations to using *C. elegans* as a model for anthelmintic screening targeting human GINs, most notably a significant false negative hit rate^31^. Thus, *C. elegans* is likely to miss some of the most promising compounds. Using a pilot study with 1,280 compounds, we devised an alternative screening model using egg-to-larva development of the free-living stages of *A. ceylanicum* human hookworm parasites, which we found predicted better anthelmintic activity against GIN adult parasites than *C. elegans* ^31^. In this work we further extended this primary screen with secondary screens using the adult stages of evolutionarily divergent hookworm and whipworm parasites and proposed an entire screening pipeline based on these results (Figure 1)^31^.

**Figure 1:**
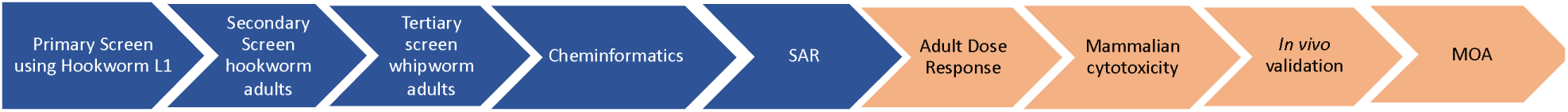
Novel anthelmintic discovery pipeline for human GINs. Our pipeline is centered around the zoonotic hookworm, *A. ceylanicum,* as a primary (larval) and secondary (adult) screening model, followed by tertiary screening against the murine whipworm *Trichuris muris* (adult) to identify broadly active compounds. Hits are subjected to deep data mining, and analogs of potential candidates are tested to generate the Structure-Activity Relationships. This report covers the results from the first five stages of the screening pipeline (in blue).

Here, we scale up this anthelmintic discovery pipeline and screen 38,293 conditions (76,586 in duplicate) representing 30,238 unique compounds. We identified 55 small molecules with *in vitro* efficacy against adult hookworms and whipworms. We focused on one novel scaffold and performed Structure-Activity Relationship (SAR) studies. Based on analogies from other systems, we propose essential GIN targets for further studies.

## Results

In our pilot screen^31^, we found that only ∼10% (4/39) of the compounds that targeted adult parasites were found by inhibiting *A. ceylanicum* egg hatch (the other 90% were found as *A. ceylanicum* larval hits). Furthermore, of those four compounds, three were retested against larval stages and were found to effectively target that stage (the fourth was not tested). On the other hand, we found that the consistency of the screen improved starting with hatched and synchronized *A. ceylanicum* first larval (L1) stages versus starting with eggs. We therefore modified the screen to start with *A. ceylanicum* L1s. We scaled up the same primary screen to include 30,000+ compounds in 13 chemical libraries (Figure 2). These libraries encompass many structural features, including diversity scaffolds, natural product derivatives, repurposing drugs, known modes of action, and target-based libraries (Table 1). In total, we screened 30,238 unique compounds in duplicate. Some compounds were screened at multiple doses due to plate format availability, making the total number of individual conditions screened 38,293 (76,586 when considered in duplicate). All libraries were initially screened in duplicate at 10 µM. As per our pipeline, primary screen hits (in both duplicate wells) were screened at 30 µM against adult *A. ceylanicum* hookworms. Adult hookworm hits were then screened at 30 µM against adult *T. muris* whipworms. A breakdown of hits “library by library” is given below and summarized in Table 1.

**Figure 2:**
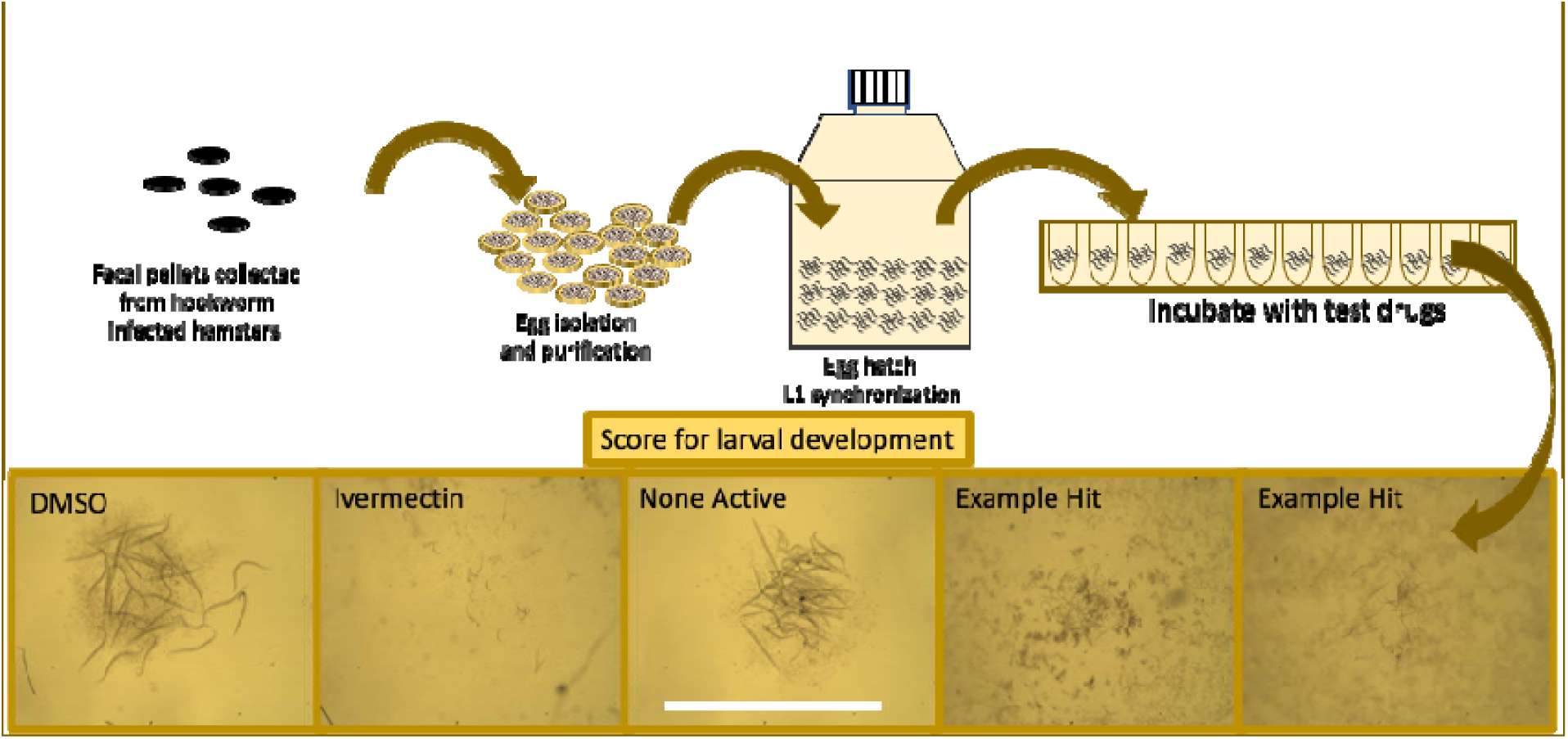
Primary screening using the free-living larval stages of *A. ceylanicum* hookworms. To work around variability in egg hatch rates, we found that starting with the first larval stage (L1) led to more consistent and robust results (see Results section). Isolated and purified eggs were allowed to hatch in S-media into synchronized L1. Each 96-well assay plate includes four wells of ivermectin (10µM) as positive control and four wells of 0.25% DMSO as negative controls. After seven days of incubation at 28°C, the plates were scored under a light microscope. A test compounds was considered active (hit) if at least 90% of all larvae in the well were killed or failed to develop. Scale bar 1 mm.

**Table 1:** Screening summary of 30,238 compounds from 13 different libraries with information on screening doses, the number of hits per screening model, and the hit rates.

| Library | Unique Compounds | Screened doses μM | Test wells screened in duplicate | *A. ceylanicum* L1 at 10μM (% hits) | *A. ceylanicum* adults 30μM (% hits) | *T. muris* Adults 30μM (% hits) |
| --- | --- | --- | --- | --- | --- | --- |
| SMSF Life Chemicals Diversity Set | 15360 | 10 | 15360 | 491 (3.2) | 33 (0.21) | 7 (0.05) |
| Broad Institute REPO 1, 2 and 3 | 6743 | 10 | 6743 | 230 (3.4) | 96 (1.42) | 36 (0.53) |
| ICCB Longwood MOA | 1245 | 10, 2, 0.4, 0.08 | 4980 | 65 (5.3) | 17 (1.36) | 9 (0.72) |
| ICCB Selleck Neuronal signaling | 1031 | 10 | 1031 | 29 (2.8) | 12 (1.16) | 2 (0.19) |
| ICCB SYNthesis Kinase Inhibitor 2 | 96 | 10, 2, 0.4 | 288 | 8 (8.3) | 1 (1.04) | 1 (1.04) |
| ICCB EMD Kinase Inhibitor 1 | 244 | 10 | 244 | 10 (4.1) | 2 (0.81) | 2 (0.81) |
| ICCB ChemBridge Focused Kinase | 250 | 10 | 250 | 2 (0.8) | 0 | 0 |
| ICCB ChemBridge Focused GPCR | 250 | 10 | 250 | 2 (0.8) | 0 | 0 |
| ICCB LINC1 Kinase inhibitors | 88 | 10, 2, 0.4, 0.08 | 352 | 4 (4.5) | 2 (2.27) | 1 (1.13) |
| ICCB LINC2 Kinase inhibitors | 88 | 10, 2, 0.4, 0.08 | 352 | 2 (2.25) | 0 | 0 |
| ICCB LINC4 Kinase inhibitors | 169 | 10, 2, 0.4, 0.08 | 676 | 8 (4.7) | 3 (1.77) | 3 (1.7) |
| SMSF Kinase Inhibitors | 492 | 10 | 492 | 13 (3) | 5 (1.01) | 3 (0.69) |
| SMSF TIMTEC Natural Compounds | 4240 | 10 | 4240 | 14 (0.33) | 0 | 0 |
| **Total** | **30238** |  | **38293** | **878 (2.9)** | **171 (0.56)** | **64 (0.21)** |

### Hits from the Life Chemicals Diversity Set

Of the 15,360 compounds in the Diversity Set tested against *A. ceylanicum* hookworm larvae, 491 were active, with a hit rate of 3.2%. These 491 larval hits were tested against the early adult stages of *A. ceylanicum* hookworm, yielding 33 hookworm adult-active compounds. An example of screening against the early adult stages of *A. ceylanicum* hookworms is shown in Figure 3. These 33 *A. ceylanicum* hookworm adult-active compounds were screened against *T. muris* adult whipworms, yielding seven hits active against the two evolutionarily divergent parasites. An example of screening against adult *T. muris* whipworms is shown in Figure 4.

**Figure 3.**
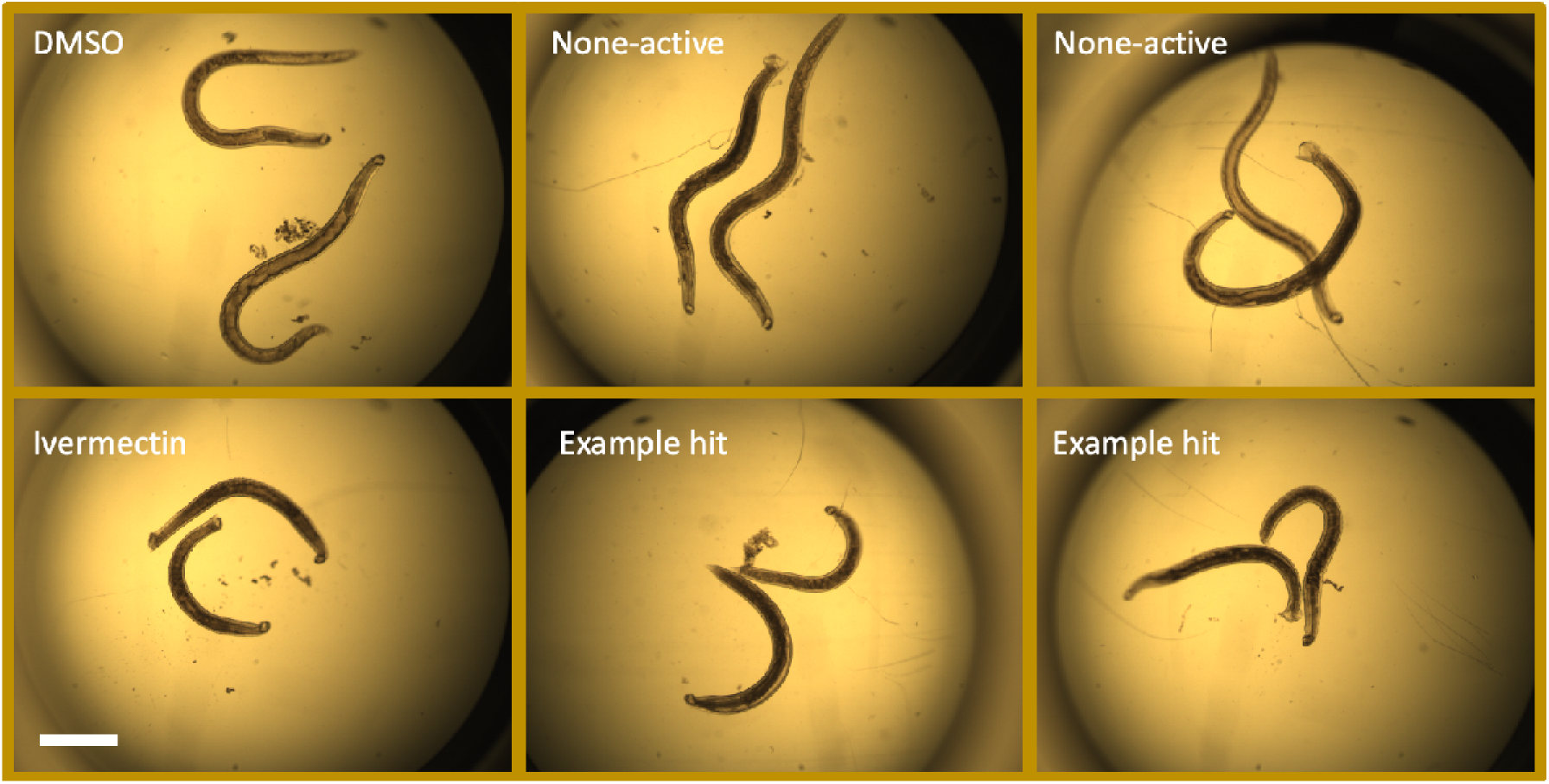
Young adult *A. ceylanicum* hookworm screens. The top panel shows healthy and motile worms from the DMSO-treated or nonactive test compounds. The lower panel shows dead or intoxicated worms from ivermectin or active compounds. Healthy worms with smooth and fully expanded cuticles were capable of controlling their body muscles, were motile, were elongated, and formed more bends. Dead or intoxicated worms were shortened, had wrinkled and shrunken cuticles, were immobile or barely mobile, and appeared more internally darkened due to damaged intestines. Because young adult hookworms are much smaller than whipworms, we manually scored these versus using the Worminator. Scale bar 1 mm.

**Figure 4:**
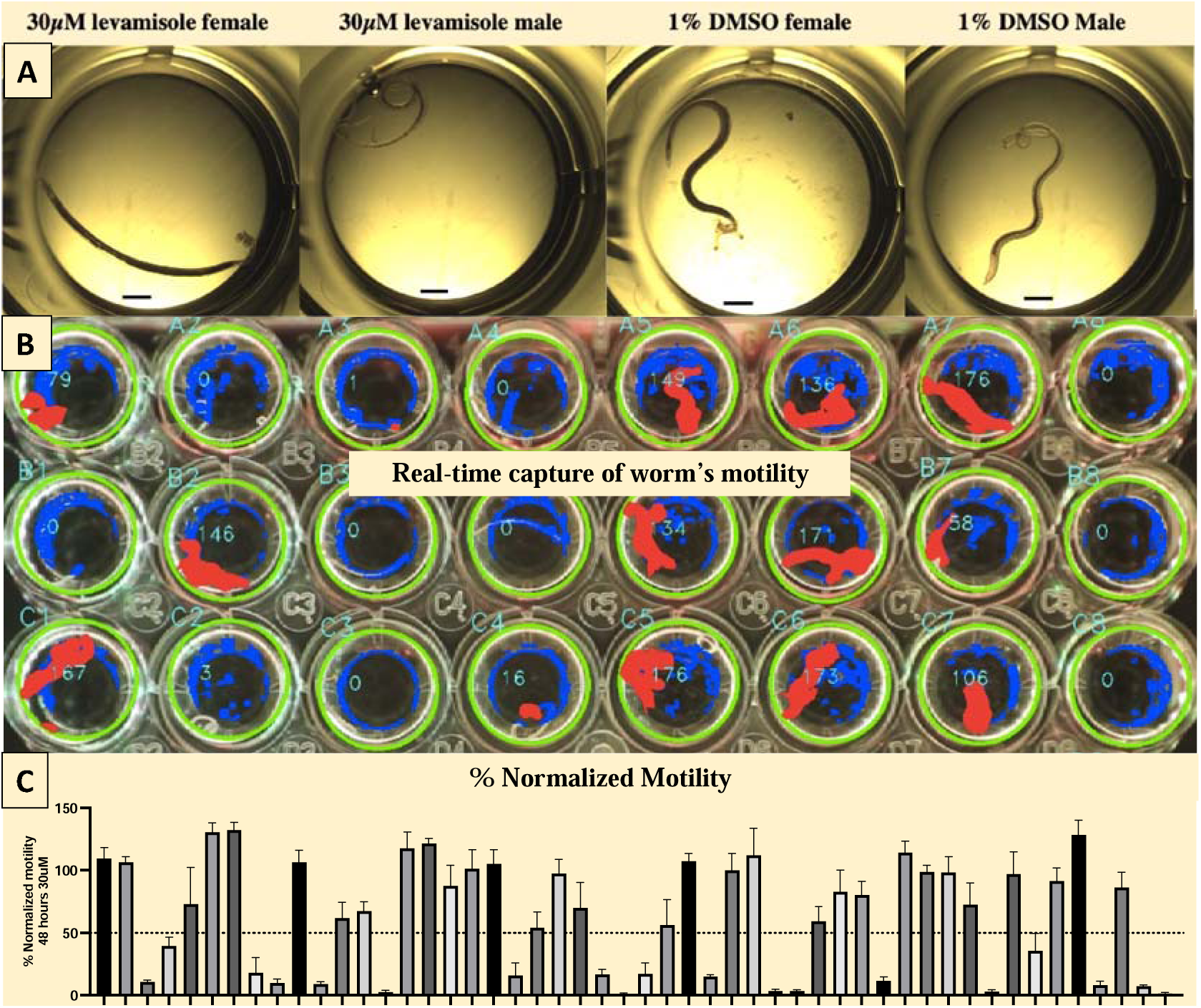
Adult parasite screening of *T. muris* whipworm. (A), Adult whipworms, male and female treated with 30µM Levamisole or 1% DMSO. Scale bar 1 mm. (B): Motility readout screenshot from 48-well plate with single adult whipworm using Wormassay-6 software and the Worminator imaging setup. Red depicts pixel changes from frame to frame (movement) and blue depicts no change in pixel (dead or very low movement). (C). Example of calculated motility readout, showing some with adult-parasitic activity with normalized motility ∼ 50% compared to the DMSO control.

### Hits from the Repurposing and Mechanism of Action (MoA) Libraries

Of the 6,743 compounds tested from the repurposing library, we identified 230 *A. ceylanicum* hookworm larval-active compounds (hit rate of 3.4%), from which 96 showed activity against adult *A. ceylanicum* hookworms (hit rate 1.42%). Of these 96 compounds, 36 were active against adult *T. muris* whipworms with a final hit rate of 0.53%. Similarly, The ICCB-Longwood MoA library contained 1,245 high-quality compounds with known mechanisms of action. We identified 65 *A. ceylanicum* hookworm larval-active compounds with a 5.2% hit rate, the highest among all tested libraries. Seventeen of these compounds carried adulticidal activity against *A. ceylanicum* hookworms, from which nine were active against *T. muris* whipworm adults, with a final hit rate of 0.72%.

### Hits from the Neuronal and the GPCR libraries

Of the 1031 compounds with known modulation of neuronal signaling, we identified 29 *A. ceylanicum* hookworm larval-active compounds (hit rate of 2.8%). Of these 29 compounds, 12 showed activity against adult *A. ceylanicum* hookworms (1.1% of the total), from which two were active against *T. muris* whipworm adults with a final hit rate of 0.19%. We also screened 250 known GPCR inhibitors and identified two compounds with larval activity against *A. ceylanicum;* however, these compounds did not show activity against the adult stages of parasites (suggesting these two compounds only target proteins essential in larval development). The general lack of success with this library could be explained by the selectivity of these GPCR inhibitors, which were tailored initially for binding to human GPCRs.

### Hits from the Kinase Inhibitor Libraries

Of seven kinase inhibitor libraries with 1427 compounds (Table 1), we identified 47 with *A. ceylanicum* larval activity, giving a hit rate of 3.2%. Of those 47 compounds, 13 showed activities against the adult stages of *A. ceylanicum* hookworms (0.9%), of which 10 compounds were also active against *T. muris* whipworm adults with a hit rate of 0.7%.

### Hits from the Natural Products Library

Of 4240 compounds from TimTec natural products libraries, we identified 14 compounds with *A. ceylanicum* larval activity, giving the lowest primary screen hit rate of 0.33%. None of these 14 larvicidal compounds showed activity against the adult stages of *A. ceylanicum* hookworms. We hypothesize that the low hit rate and lack of adult activity are due to the nature of these libraries (see Materials and Methods).

### Summary of 30,238 compound screen

Collectively, we identified 878 compounds with *A. ceylanicum* larval activity, giving an overall primary screen hit rate of 2.9% (Table 1). In our secondary screen, we tested all larval-active compounds at 30 µM against the adult stages of *A. ceylanicum* hookworm parasites. Of those, 171 (19.5% of the larval hits) had anthelmintic activity toward adult hookworms, representing a hit rate of 0.5% based on the total number of compounds in the initial libraries. These 171 hookworm-adult actives were tested against adult *T. muris* whipworms at 30 µM. Sixty-four compounds (37% of the adult hookworm hits and 0.21% based on the total number of compounds in all the libraries) showed activity against whipworm adults (Table 1). Of the 64 compounds identified from all libraries, nine were redundant (included independently in multiple libraries), yielding 55 unique active compounds showing a motility index of ≤1 or intoxicated phenotype in hookworms and relative motility of ∼ 50% in whipworms compared to the DMSO control, listed in Table 2.

**Table 2:**
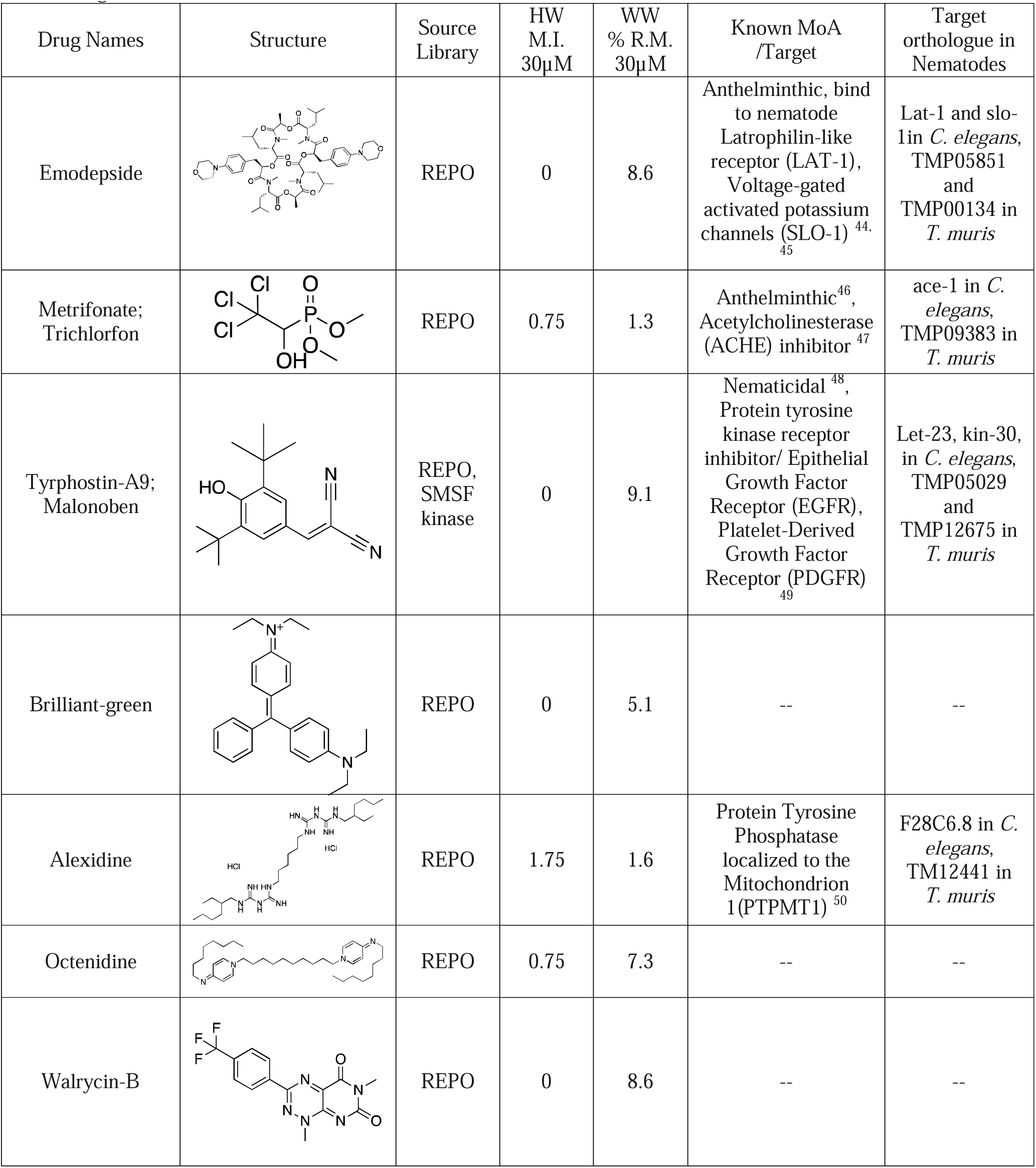
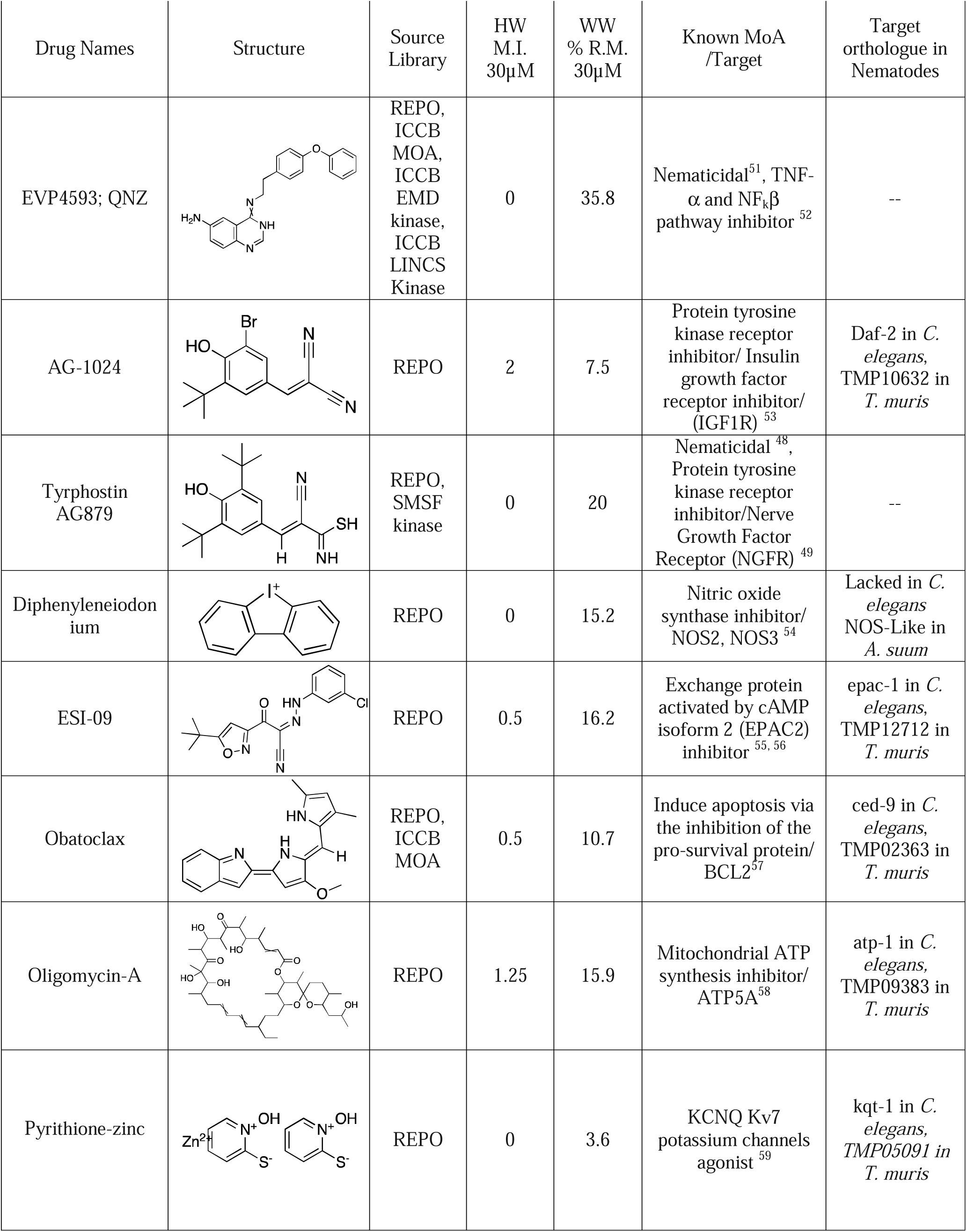
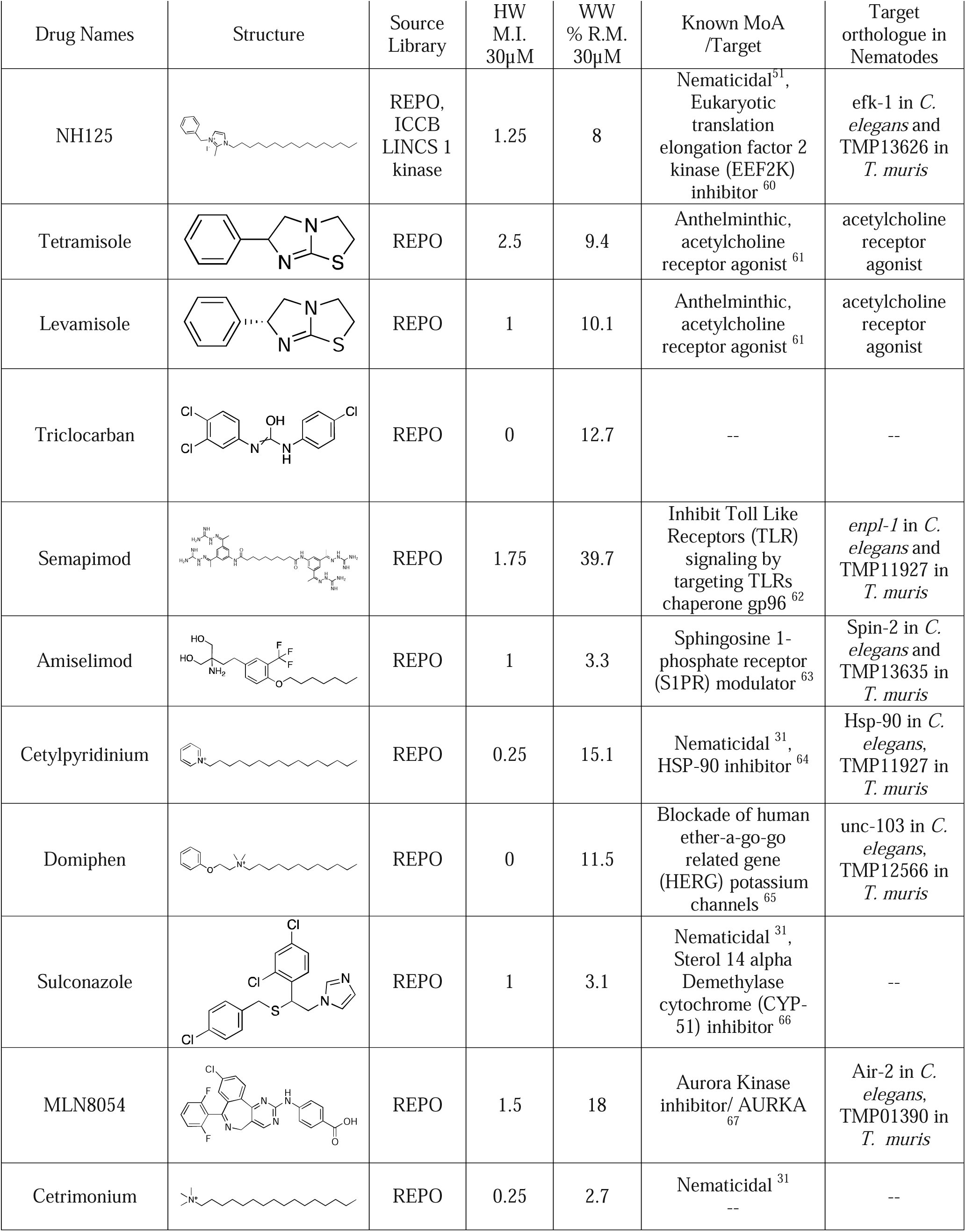
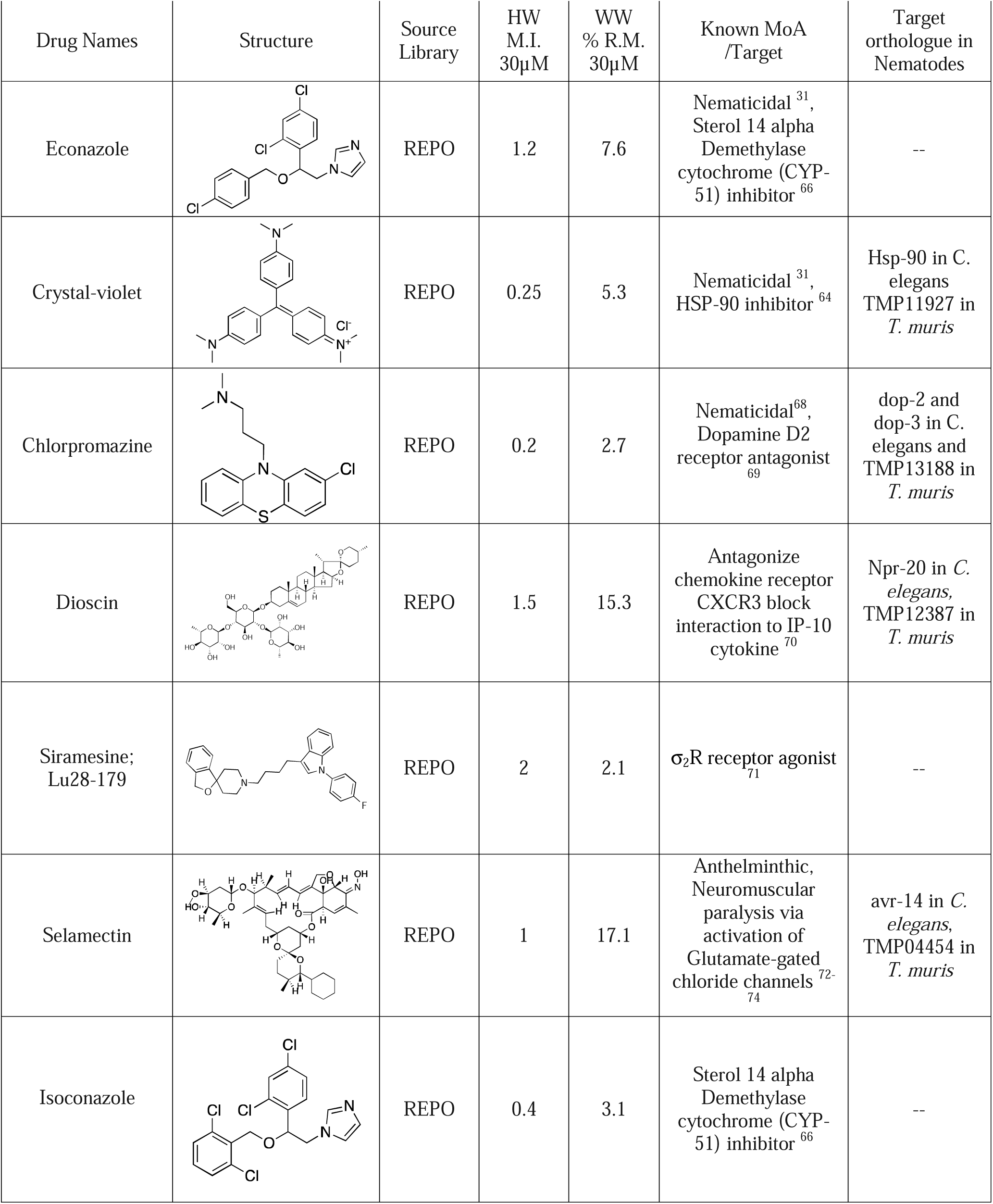
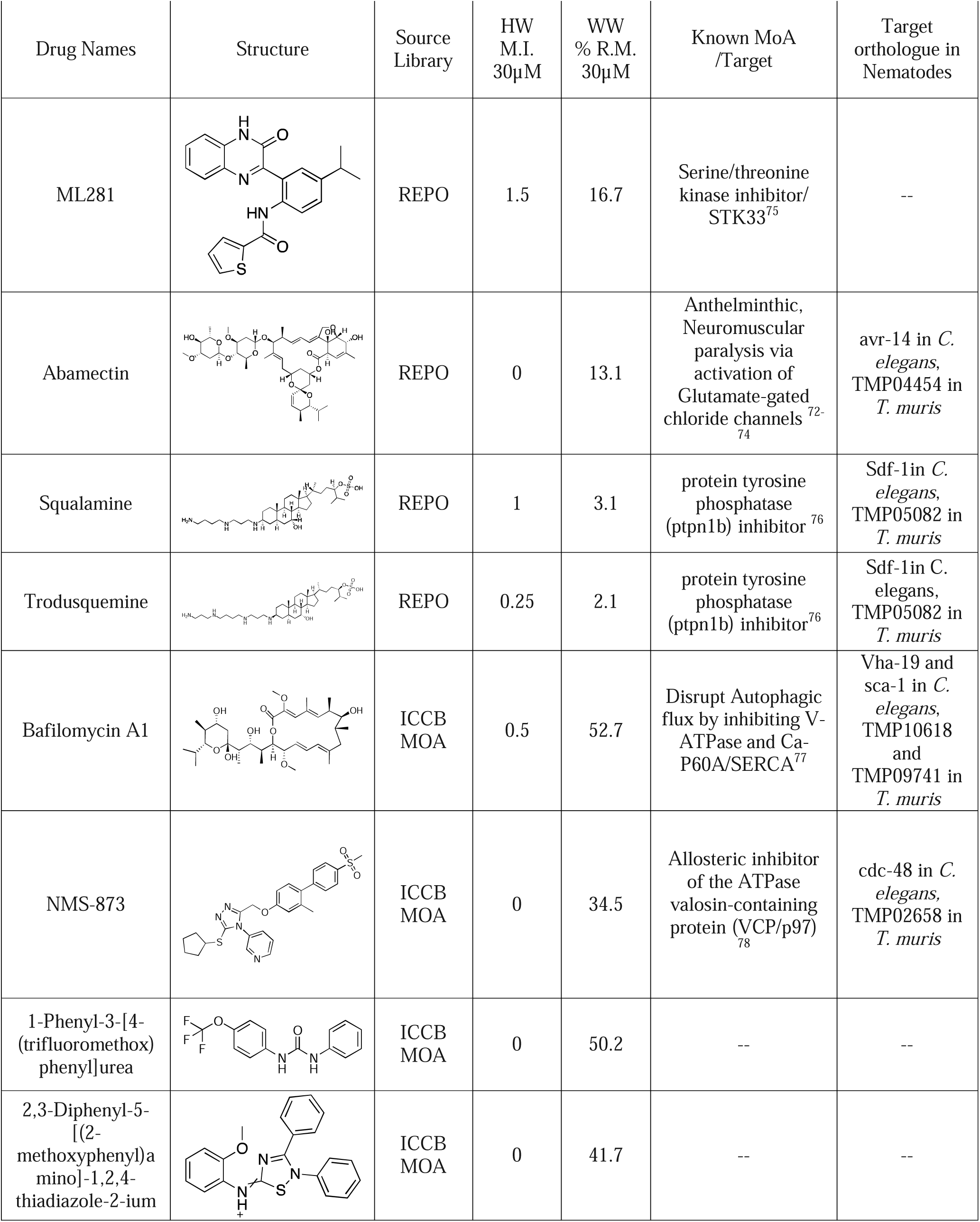
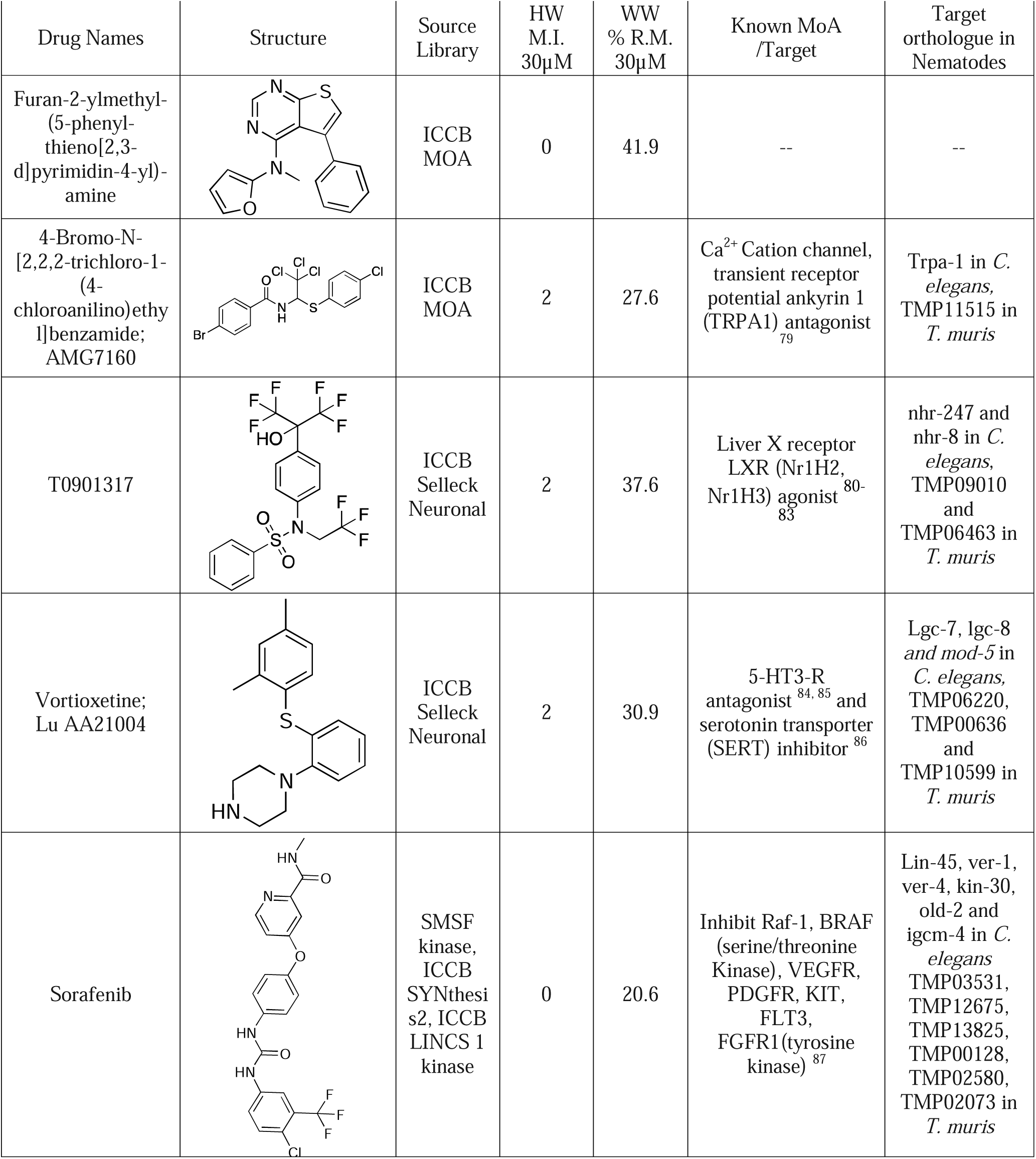
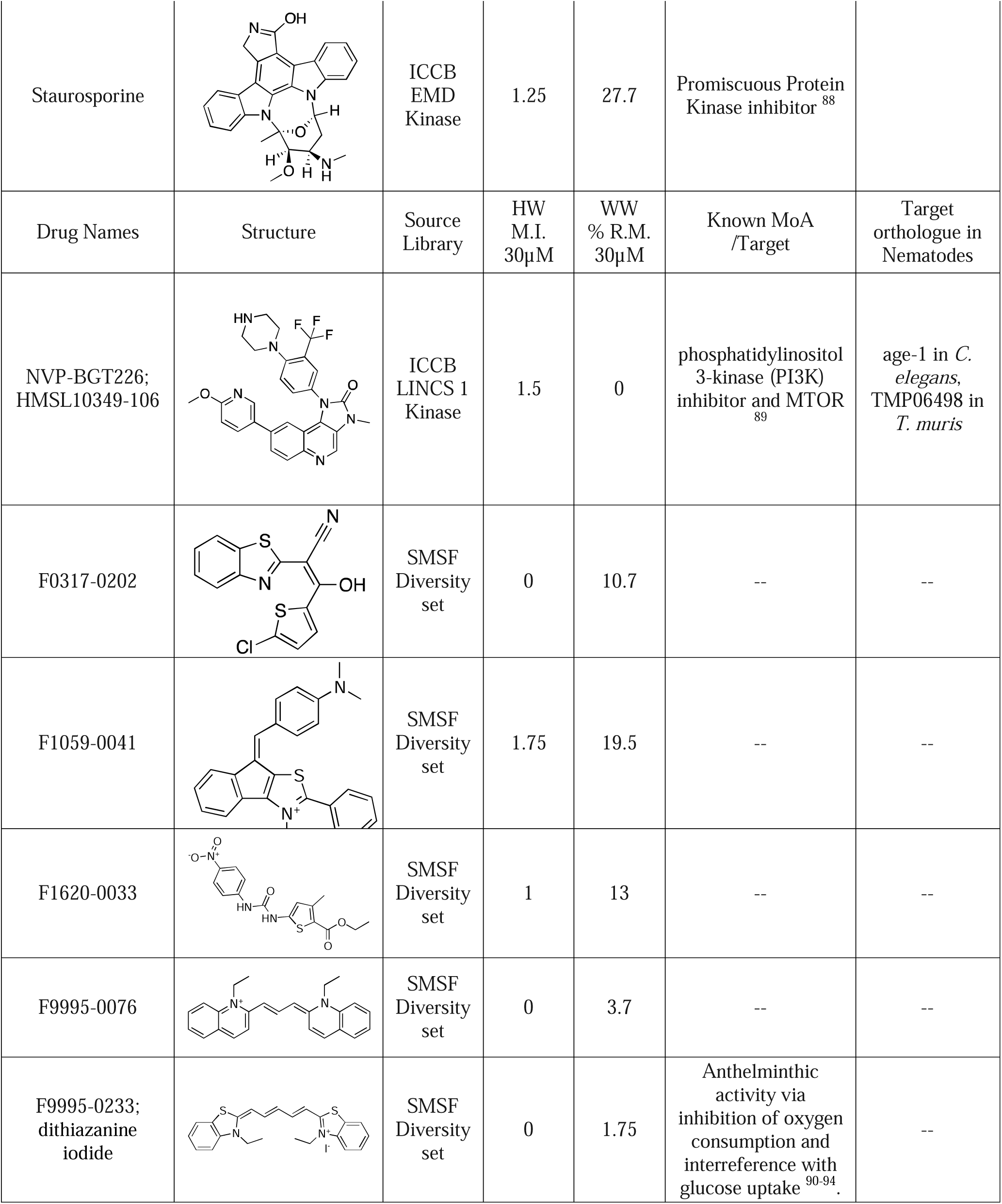
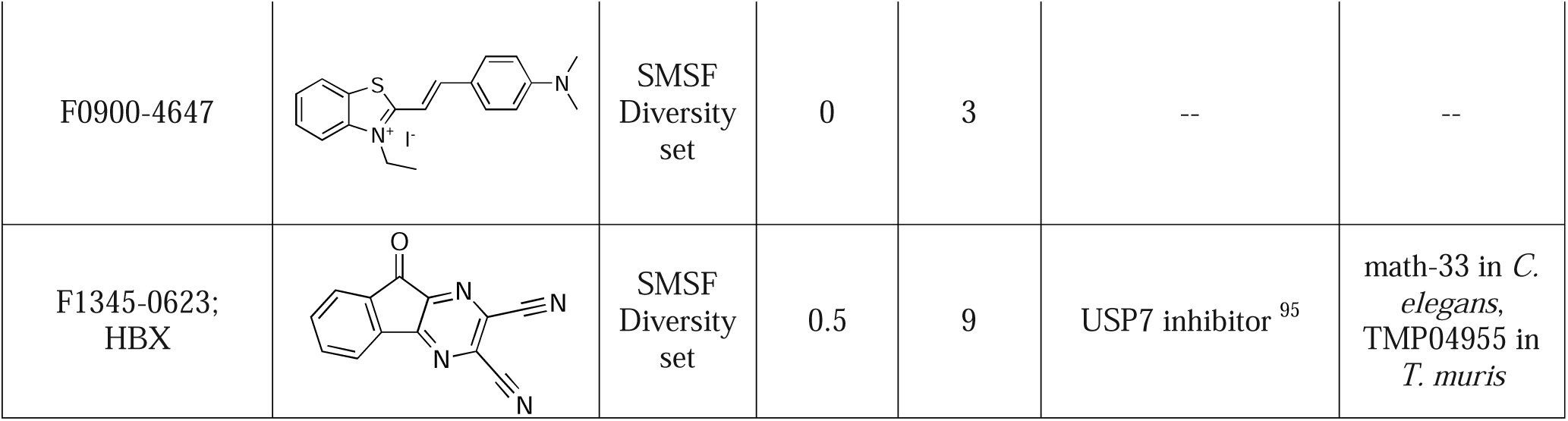
List of 55 potent and broadly active compounds including chemical structure, source library, *in vitro* activity against *A. ceylanicum* hookworms (HW) with motility index (M.I.) of ≤1 or intoxicated phenotype, where 1 is intoxicated worms moving only aftet being touched and 0 is completely immotile), *in vitro* activity against *T. muris* whipworms (WW) with percent Relative Motility (% R.M.) of ∼ 50%, where100% is healthy; 0% is non-motile) to the DMSO control, known mode of action in eukaryotic organisms, and the orthologous targets in nematodes if any. (--) no known target.

Screened libraries contained known anthelmintic compounds, and we successfully identified albendazole, oxibendazole, flubendazole, methiazole, oxfendazole, thiabendazole, ivermectin, abamectin, dormamectin, emamectine, eprinomectin, selamectin, emodepside, closantel, monepantel, and moxidectin as part of the primary and secondary *A. ceylanicum* hookworm screens. However, many of these anthelmintics didn’t show activity in our tertiary screening against the *T. muris* whipworms at 30µM, highlighting the whipworms’ resilience to anthelmintics compared to hookworms. For example, the published 50% inhibitory concentrations (IC_50_) of albendazole and ivermectin against *T. muris* whipworms are both > 200µg/ml^43^. Only anthelmintic compounds that showed activity at 30µM against *A. ceylanicum* and *T. muris* are shown in Table 2.

### Data Mining

The 55 broadly active compounds were subjected to a literature search to identify, if known, their possible mode of action and the eukaryotic drug targets in other organisms and the orthologous targets in phylum Nematoda, focusing on *C. elegans* in clade V and *T. muris* in clade I (Table 2). The compounds encompass a wide range of putative target classes including potassium channel, glutamate-gated chloride channel, acetylcholinesterase, acetylcholine receptor, tyrosine kinases, aurora kinase, phosphatidylinositol 3-kinase (PI3K), tyrosine phosphatase, insulin growth factor receptor (IGF1-R), nitric oxide synthase (NOS), Exchange Protein Activated by Cyclic AMP (EPAC), pro-survival protein (BCL2), ATP synthase, translation elongation factor, Enoyl-acyl carrier protein reductase (ENR), p38 mitogen-activated protein Kinase (p38 MAPK), sphingosine 1-phosphate receptor (S1PR), Heat Shock Protein-90 (HSP-90), Dopamine D2 receptor, liver X receptor (LXR), serotonin (5-HT)3 receptor, chemokine CXCR3 receptor, and ubiquitin specific peptidase 7 (USP-7) and Phosphatase localized to Mitochondrion 1(PTPMT1). Compounds identified here with well-known targets and mechanisms of action in other organisms may be used as probes for druggable and essential ortholog targets in nematode parasites (Table 2). For example, alexidine was shown to induce selective responses similar to the knockdown of its target PTPMT1^96^. Similarly, T0901317 is a well-known selective agonist of LXR used in many studies to investigate the physiological effects of LXR on cholesterol and lipid homeostasis; notably, T0901317 induction of LXR was found to reduce cholesterol biosynthesis and to induce cholesterol breakdown into bile acids ^80–82^. Nematodes can’t synthesize cholesterol and rely on exogenous sources from the environment or their hosts^97, 98^. Thus, ligand activation of the LXRs ortholog in parasitic nematodes could result in cholesterol depletion and negatively impact parasite lipid homeostasis. As examples, we discuss putative nematode targets of four of these--BCL2, NOS, PTPMT1, and LXRs, selected because these targets are known to be essential (See Supplementary Discussion).

### Novel anthelmintic scaffold

We identified seven compounds with broad anthelmintic activity from the Life Chemicals Diversity Set, which is compiled of *de novo* and newly synthesized molecules developed for initial phenotypic screening. Because of the novelty of this library, we chose to study one of the hits further as an initial foray into follow up studies. Among the seven actives from this library of novel scaffolds, we focused on F0317-0202 as a lead for further studies because given its better drug-like structure over the other six (and see Discussion). To further investigate, 28 commercially available analogs of F0317-0202 were identified using structural similarity searches and then purchased from Life Chemicals. These could serve as training sets for ligand-based optimization and generate the SAR model. We screened this set of analogs against adult *T. muris* whipworms, followed by older and larger *A. ceylanicum* hookworm adults, which could be read more easily in the Worminator.

Of the 28 analogs, we identified 12 compounds (41%) that inhibit *T. muris* whipworms motility by ≥ 50% relative to the DMSO control (Figure S1). All 12 were tested and found to be active against *A. ceylanicum* hookworm adults (Figure S1). To rank these 12 analogs, they were tested *in vitro* in dose-response against hookworms and whipworms. Of these 12 broadly active compounds, six analogs (F0317-0161, F0317-0013, F0328-0210, and F0317-0009, F0317-0019, and F0317-0160), along with the parent, showed IC_50_s of ≤5µM against adult hookworms and ≤25µM against adult whipworms (Figure 5). Note for comparison, the IC_50_ values of albendazole and ivermectin, the first-line anthelmintics, against *T. muris* whipworms were > 753 and 228µM, respectively ^43^.

**Figure 5:**
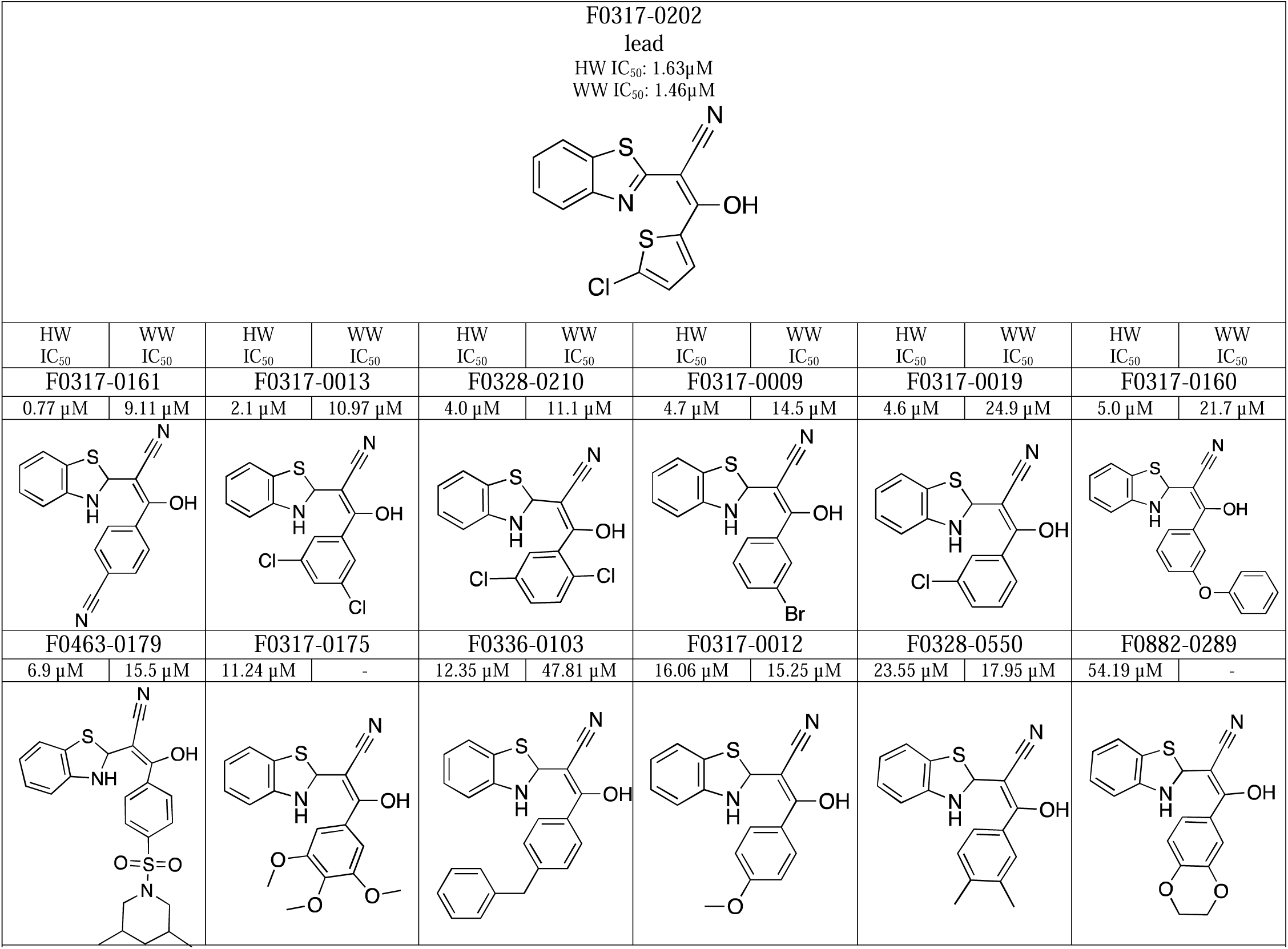
IC_50_values of 12 F0317-0202 broadly active analogs: Chemical structure of the top tested analogs and the motility IC_50_values, against *A. ceylanicum* hookworms (HW) and *T. muris* whipworms (WW). (-) indicate that we were not able to calculate an IC_50_ values as the highest dose used did not reach 50% inhibition.

Based on these data, an F0317-0202 initial SAR model for potent and broad anthelmintic activity was built and found to be quite strict— slight modifications could abolish the anthelmintic activity entirely. The functional core identified in this potent scaffold was the (z)-2-(2,3-dihydrobenzo[*d*]thiazol-2-yl)-3-hydroxybut-2-enenitrile (Figure 6A) linked to an aromatic ring, either thiophene or benzene (Figure 6. B). Interestingly, replacing thiophene with benzene resulted in a dramatic reduction in the activity against adult whipworms. Analogs with a benzene ring and one or more of the following groups: nitrile, ethylbenzene, chlorine, bromine, or anisole (Figure 6. C) showed potent *in vitro* activity against both parasites (Figure 5). F-0317-0161, with a benzonitrile, showed the most potent *in vitro* activity against hookworms with an IC_50_ of 0.77µM. These findings suggest that further optimization and finetuning on the variable aromatic ring is feasible to design analogs with broad-spectrum anthelmintic activity.

**Figure 6:**
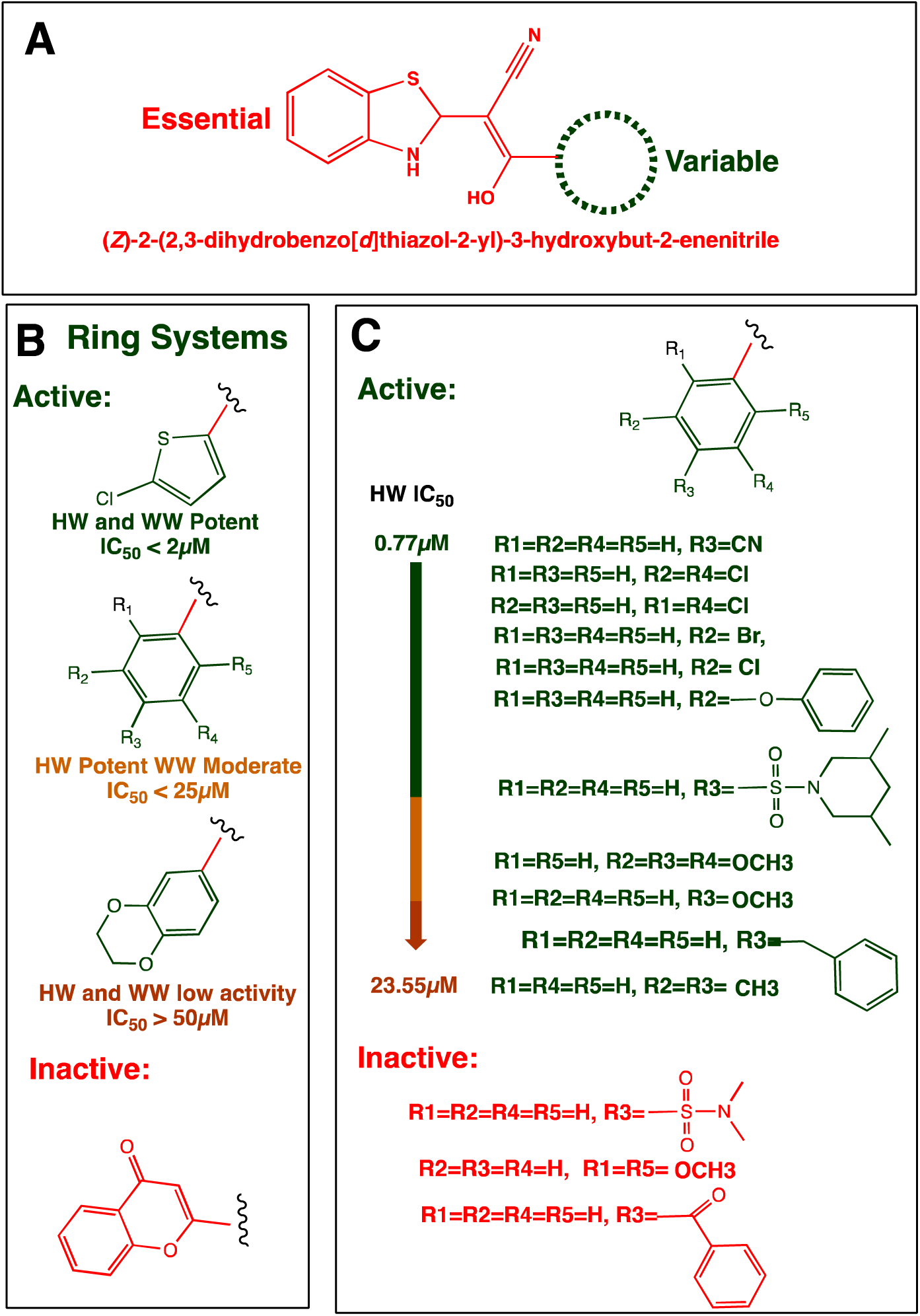
F0317-0202 Structure-activity relationship (SAR) model. (A) The core and essential structure (red) and the position of the variable rings of F0317-0202. (B) variable rings and the associate nematicidal activity. (C) Analogs with a benzene ring and variable groups ranked based on the IC_50_ against the adult stages of hookworms.

## Discussion

Human GINs are among the most prevalent infections of the NTDs. Because understanding of these parasites’ basic biology is still limited, whole organism screening (*i.e.*, phenotypic screening vs. target-based screening) remains the most effective approach in finding effective anthelmintics. We previously validated a novel screening pipeline using a set of 1280 FDA-approved compounds from a single library ^31^. This screening pipeline utilizes two phylogenetically diverse nematode parasites, *A. ceylanicum* hookworms and *T. muris* whipworms, located at opposite ends of the phylum Nematoda phylogenetic tree and that are either a human parasite (*A. ceylanicum*) or closely related to a human parasite (*T. muris* vs *T. trichiura*). By selecting these diverse nematodes, we aim to identify compounds with broad anthelmintic activity. Here, using much larger and more diverse sets of compound libraries, we used our pipeline to screen more than 30,000 compounds, which is unprecedented for targeting the GINs of humans.

The life chemicals diversity set library comprises 15,360 compounds representing 1,717 chemical scaffolds from novel and newly synthesized small molecules. Using our screening pipeline, we uncovered seven molecules with broad activity against parasitic nematodes, giving a hit rate of 0.045%. Among these seven, four are polymethine cyanine dyes: F1059-0041, F9995-0076, F0900-4647, and dithiazanine iodide (F9995-0233). Dithiazanine iodide is known to be active against hookworms, threadworms, roundworms, pinworms, and whipworms ^90–94^, providing additional validation for our pipeline. Of the other three non-polymethine cyanine dyes, F1345-0623 has been identified as a potential inhibitor (IC_50_ in the sub-micromolar range) of the deubiquitinase USP7 ^95^; homologous proteins have been found to play essential roles in *C. elegans* (math-33)^99^ and *Schistosoma mansoni* (SmUSP-7) ^100^.

Of the remaining two novel compounds, F0317-0202 showed better IC_50_s than F01620-0033 against hookworms (1.6 compared to 14µM) and whipworms (1.4 compared to 5.8µM) and better drug-like structure and thus was selected for further studies. F0317-0202 includes two aromatic ring systems (thiophene and benzothiazole), listed among the most frequently used ring systems from small molecules in the FDA Orange Book ^101^. F0317-0202 also has two functional groups (OH and CN), listed among the most frequent functional groups occurring in bioactive molecules ^102^. F0317-0202’s physicochemical property better agrees with Lipinski’s rule of five for drug-likeness than F01620-0033^103^, with a molecular weight of 318.8 compared to 449g/mol, one hydrogen bond donor compared to two, five hydrogen bond acceptor compared to seven. Finally, F0317-0202 was predicted to have better absorption with 113□^2^ of Topological Polar Surface Area (TPSA) than F01620-0033 (142□^2^), as compounds with smaller TPSA are predicted to have better absorption ^104^. Out of 28 similar analogs, 12 compounds (42%) showed potent activity against whipworm and hookworm parasites, with multiple compounds showing promising activity against the two parasites. We consider F0317-0202 and at least six of its analogs, F0317-0161, F0317-0013, F0328-0210, and F0317-0009, F0317-0019, and F0317-0160, as well-positioned for further characterization, mode of action studies, lead optimization, cytotoxicity studies in mammalian cells, and *in vivo* validation.

In contrast to screening diversity sets or random compound collections, we also screened repurposing libraries, which contain compounds already approved or associated with clinical trials and which can serve as a source for novel therapeutics for new indications ^105^. The Broad Institute repurposing library, which we used in our screen, contains 6,743 compounds from molecules with high structural diversity and with 2006 known protein targets, including but not limited to the G protein-coupled receptors (GPCR), acetylcholine receptors, protein kinases, oxygenases, voltage-gated channels, and nuclear hormone receptors^106^. In addition, screening compounds with known targets can be highly beneficial, especially against NTDs where a deep understanding of their biology is still lacking. Compounds with known modes of action offer a significant advantage in drug development because they can act as probes for essential targets, assuming the targets are known. For this reason, we also screened the ICCB-Longwood mechanism of action library containing 1,245 high-quality compounds with known targets.

The total number of compounds tested from both libraries was 7,988, uncovering 45 compounds with putative broad activity against parasitic nematodes, giving a hit rate of 0.56%, 12.4 times that obtained from the generic diversity set library. These results are the first to thoroughly compare a generic dissimilarity-based library type with repurposing/known mode of action compounds against parasitic nematodes using the same screening conditions and pipeline. Our results indicate that repurposing/known mode of action libraries are significantly more (>10X; P< 0.00001 Fisher’s exact test) likely to uncover active compounds.

The plethora of the chemical universe is expected to contain 10^12^ to 10^180^ drug-like compounds^107^, which are impossible to screen against any single organism, not to mention GINs. Therefore, techniques and strategies to design chemical libraries covering essential pharmacological targets are highly recommended. Among these approaches is the compilation of a target-focused library of compounds, which can be fitted by molecular docking to bind and inhibit the function of a specific target or protein family^104, 108^. This strategy has been previously validated using a kinase-targeted library of 1440 compounds tailored to fit in a total of 41 kinases from five different families, showing a 6.7-fold increase in the overall hits compared to the generic collection of molecules^109^. Target-focused screening improves efficiency by excluding compounds unlikely to modulate known target proteins of interest and is associated with increased potency and specificity^110^.

We found that target-based libraries could serve as good sources for anthelmintic discovery. We screened nine target-focused libraries, including one for neuronal targets, one for GPCRs, and seven libraries targeting protein kinases. Neuronal signaling and the neuromuscular system are essential for nematodes’ survival, including feeding, mating, egg laying, and locomotion; thus, many anthelmintics drugs target the nematode nervous system^111^. Neuropeptide GPCRs have been identified as potential and druggable targets and were found to play critical roles in the helminth neuromuscular function^112^. Kinases are the second most targeted protein class after GPCRs, with over 200 kinase inhibitors having reached the clinical phases and 52 FDA-approved drugs targeting protein kinases^39, 113^.

Together, these nine libraries contain 2,708 compounds, and within these, we identified 12 compounds with putative broad activity, giving a hit rate of 0.44%, statistically the same as that obtained from the combined repurposing and mode of action libraries (P=0.54, Fisher’s exact test). It is worth noting that three of the nine target-based libraries resulted in no hits with activity against hookworms and whipworms: the ChemBridge Focused kinase, GPCR, and the LINCS 2 kinase inhibitors, which could be explained by the lack of specificity to the parasitic kinases and GPCRs, as these libraries were fitted to selectively binding specific human targets.

Historically, natural products have been the primary source of remedies for human ailments, and purified natural molecules or direct derivatives have contributed to at least 34% of all drugs approved by the FDA between 1981 and 2010^114^. Using our novel pipeline, we screened TimTec libraries of 4,240 purified compounds derived from or synthetically based upon natural products (see Materials and Methods). We identified 14 compounds with *A. ceylanicum* larval activity. However, none of these larvicidal compounds carried activity against adult parasites. This result may reflect biases of these particular libraries (see Materials and Methods).

In summary, we have developed a novel high throughput screening pipeline to identify anthelmintic compounds for GIN parasites and screened 13 different libraries with a total of 30,238 compounds that led to the discovery of 55 compounds with potent activity against two species of evolutionary distant nematode parasites. Repurposing, mode of action, and target-based libraries were more profitable than the diversity set for anthelmintic screening. For these 55 compounds, we reported the known modes of action and related targets in other organisms, and potential counterparts in the nematode, if available. As an example, we highlighted four nematode targets as potential candidates for target-based drug screening. We also identified thiazolyl oxopropanenitrile as a novel chemical scaffold with broader anthelmintic activity identified from the diversity set library and established the initial SAR models for potent activity against hookworms and whipworms. Our findings open the doors for multiple follow-up approaches, including target identification, target-based screening, and ligand or structure-based pharmacophore optimization for developing safe and parasite-selective molecules.

## Materials and Methods

### Ethics statement

All experiments involving animals were conducted per the recommendation of the National Institute of Health (USA) Guide for Care and Use of Laboratory Animals and the Animal Welfare Act. The animal protocols used in this study (PROTO202000044 and PROTO202000071) were approved by the Institutional Animal Care and Use Committee (IACUC) of UMASS Chan Medical School.

### Parasite maintenance in laboratory animals

Following standard protocols, *A. ceylanicum* hookworm parasites were maintained in golden Syrian hamsters^115^, and *T. muris* whipworm parasites were maintained in STAT6-/- mice ^116^.

### Primary screening using Larval Developmental Assay

Feces from *A. ceylanicum-*infected hamsters were collected ∼18 days post-infection and about 20 grams of feces were processed for egg isolation and purification following salt/sugar flotation protocol^117, 118^. Purified and surface sterilized eggs were incubated in S-medium^119^ at 28 °C, without any food source for about 20 hours, allowing eggs to hatch into synchronized first-stage larvae ^118^. Newly hatched L1 were washed three times in a S-medium. About 30-35 L1 larvae were dispensed into each well of 96-well assay plates containing 90µL of S-media supplemented with 100U/ml of penicillin, 100µg/ml streptomycin, and 2.5µg/ml amphotericin, *Escherichia coli* OP50 with final OD600 of 0.3 and 0.15µl Dimethylsulfoxide (DMSO). Library plates were diluted in S-media to 100µM and 1%DMSO. Using a multichannel pipette, 10µL of the library plates were transferred into each well of the assay plates, giving a final concentration of 10μM of test drug and 0.25% DMSO. Plates were prepared in duplicates. Test plates were incubated at 28 °C for seven days and then scored for development under a dissecting microscope. Test drugs were considered hit if > 90% of L1 larvae failed to develop in duplicate wells. Each assay plate includes four wells of 10µM ivermectin as positive control and four wells of 0.25% DMSO as negative control. For a group of 12 hamsters collecting feces twice weekly, the throughput for this assay is ∼2200 compounds/week in duplicate. Cherry-picked compounds for follow up study (secondary screen), were obtained from the original library sources (Broad, Harvard, UMASS Chan Med) independent of the initial library plates given to us for the primary screen.

### Secondary screening using young adults of hookworm parasites

We next screened hits from the primary screening (larval development) against the adult stages of hookworm parasites. *A. ceylanicum* hookworm young adults were harvested 10-11 days post-inoculation from the small intestines of infected hamsters. Harvested worms were washed and incubated in prewarmed hookworm culture medium (RPMI1640 of pH 7.2 supplemented with 10% fetal bovine serum, 100U penicillin, 100μg/ml streptomycin, 2.5μg/ml amphotericin). Two adults of *A. ceylanicum* hookworm were manually picked into each well of the 96-well assay plates, containing 70µL of hookworm medium supplemented with 0.7µl DMSO. Using a multichannel pipette, 30µL from the library plates diluted in hookworm medium to a final concentration of 100µM and 1% DMSO were dispensed into screening wells, giving a final concentration in assay plates of 30μM of test compounds and 1% DMSO with duplicate wells/drug. Assay plates were incubated for 48 hours at 37 °C and 5% CO_2_. Drug activity was determined using the standard motility index ^31, 116^. A motility index of 3 was given to vigorous healthy adults, 2 for motile but slow adults, 1 for immotile adults that moved after stimulation by touching, and 0 for non-motile adults even after touching. The drug was considered a hit if the motility index of at least three of the four *A. ceylanicum* hookworm adults were scored ≤1 and/or if all the parasites in the well had significant morphological defects (*e.g.*, shrunken cuticles, darkened intestines (Figure 3). Each assay plate included four positive control wells (30µM ivermectin) and four negative control wells (1% DMSO). Cherry-picked compounds for follow up study (tertiary screen), were obtained from the original library sources (Broad, Harvard, UMASS Chan Med) independent of the library plates given to us for the primary and secondary screens.

### Tertiary screening using adult whipworm parasites

We next screened *A. ceylanicum* hookworm adult hits against *T. muris* whipworm parasites. *T. muris* adult whipworms were harvested from the cecum and large intestines of infected STAT6 -/- mice. Single worms were manually picked into each well of a 48-well plate, six wells/ test drug, and a final volume of 291µL of whipworm culture medium with 2.1µl DMSO (like hookworm culture medium but supplemented with 5% fetal bovine serum). 9µL from library plates diluted in whipworm medium to a final concentration of 1mM and 10% DMSO were dispensed into each well of the screening plates, giving a final concentration of 30μM test drug and 1% DMSO. Assay plates were incubated at 37 °C, and 5% CO_2._ *T. muris* whipworm motility was measured using an in-house assembled worminator and worm assay software^120^ for 3 minutes after 24 and 48 hours. Drug activity was determined using the percent of relative motility compared to DMSO control. Drugs were considered active if the 24- and/or 48-hour average motility of six wells (1 worm/well) was reduced by at least 50% compared to negative control (Figure 4).

### Chemical Libraries Screened

Libraries are provided by the Broad Institute (REPO), Harvard Medical School (ICCB), and UMASS Chan Small Molecules Screening Facility (SMSF), all of which have the purity and identity of their libraries assured by the source company or in-house using liquid chromatography-mass spectrometry.

### Life Chemicals 15K diversity set

Life Chemical pre-plated diversity set library was obtained from SMSF, UMASS Chan Medical School. The library contains 15,360 compounds representing 1,717 chemical scaffolds with an average Tanimoto similarity index of 0.409. Library compounds provided in 96-well plates with 10 mM stock concentration in 100% DMSO. Hits from the primary and secondary screening were cherrypicked into 96-well plates for *A. ceylanicum* hookworms or *T. muris* whipworms screening.

### Broad Institute Repurposing Hub

The drug repurposing hub includes 6,743 compounds which were from a curated and annotated collection of preclinical drugs in phase 1 (REPO 1) or phases 2-3 (REPO 2) and FDA-approved and launched drugs (REPO 3). REPO drugs cover a wide range of disease areas, including but not limited to infectious diseases, allergies, cardiology, dermatology, endocrinology, metabolism, neurology, and gastroenterology. The library covers 1,133 different modes of action and 2,183 different drug targets^106^. Library compounds provided in 96-well plates with 10 mM stock concentration in 100% DMSO. Hits from the primary and secondary screening were cherrypicked from the source for *A. ceylanicum* hookworms or *T. muris* whipworms screening.

### ICCB Screening libraries

The Institute of Chemistry and Cell Biology-Longwood (ICCB-L) Screening Program at Harvard Medical School offers a wide range of well-annotated drug screening libraries. We have screened multiple ICCB-L libraries, including mechanism of action, neuronal targets, GPCR-based inhibitors, and six libraries of kinase inhibitors.

*ICCB-L MOA*: the mechanism of the action library is a collection of high-quality annotated compounds with biological activity against known targets with target-compound pairs based on high target specificity and potency. The library contains 1,245 unique compounds, each plated in 4 doses (10, 2, 0.4, 0.08 mM) in 100% DMSO. *ICCB-L Selleck Neuronal Targets:* the neuronal targets library was initially composed of Selleck chemicals from FDA-approved compound sets that were determined to have biological activity in neurologic research and neurological assays. Library compounds were provided in 96-well plates with 10mM stock concentration in 100% DMSO.*ICCB-L ChemBridge focused GPCR:* This focused library of 250 GPCR inhibitors was selected from 13,000 GPCR inhibitors assembled by ChemBridge. The library has 11 different scaffolds, each containing 19-24 compounds. Library compounds were provided in 96-well plates with 10mM stock concentration in 100% DMSO.

*Kinase inhibitor libraries:* in total, we screened focused sets of 1428 kinase inhibitors from 7 different kinase libraries, some of which were screened at multiple doses.

*ICCB-L SYNthesis kinase inhibitor:* this library was assembled by SYNthesis Med Chem of 96 compounds known to inhibit different kinases, each plated in 3 doses (10, 2, 0.4 mM) in 100%DMSO. *ICCB-L EMD Kinase inhibitors:* this library was built by EMD of 244 protein kinase inhibitors. Library compounds were provided in 96-well plates with 10mM stock concentration in 100% DMSO. *ICCB-L ChemBridge Focused Kinase inhibitors:* This library of 250 compounds was created and selected from the ChemBridge kinase-biased library of 6000 compounds. These compounds were selected based on the pharmacophore properties of known kinase inhibitors or from compounds predicted to interact with the ATP binding site of kinases. In total, these compounds represent 16 different scaffolds that cover 34 kinase targets. Library compounds were provided in 96-well plates with 10mM stock concentration in 100% DMSO. *ICCB-L LINCS 1,2 and 4 Kinase inhibitors:* these three libraries were assembled by LINCS together, including 345 compounds with known or predicted kinase inhibitors, each plated in 4 doses (10, 2, 0.4, and 0.08 mM) in 100%DMSO.

### SMSF kinase inhibitor library

This library includes 429 kinase inhibitors with activity and safety confirmed by preclinical research and clinical trials. Library compounds were provided in 96-well plates with 10 mM stock concentration in 100% DMSO.

### Natural Products

we screened the TimTec library of 4240 purified compounds from natural products or derivatives. Library compounds were provided in 96-well plates with 10 mM stock concentration in 100% DMSO. The TimTec library was obtained from the SMSF and includes two types of libraries with no overlap between them: (1) the Natural Products Library (NPL), which contains compounds are primarily sourced from plants with the remaining samples from bacteria, fungus, and animal source; and (2) the Natural Derivatives Library (NDL), which is a hybrid between pure natural molecules and synthetic organic chemistry, derived from publications and TimTec in-house material. The NDL elaborates on structural diversity of pure natural compounds including natural derivatives, analogs, semi-natural compounds, and mimics.

### Structure-Activity Relationship (SAR) of anthelmintic hits

Using vendor databases and online tools, 28 analogs of F0317-0202 were identified and purchased from Life Chemicals. Each analog has slight structural modification compared to the parent molecule. These analogs were tested against the adult stages of *T. muris* whipworm and *A. ceylanicum* hookworm parasites at 30µM and scored using the Worminator at 24 and 48 hours. Hence, *A. ceylanicum* hookworm parasites used in testing F0317-0202 analogs for SAR model were mature adults harvested 18 days post-infection.

### Dose-response against hookworms and whipworms

To better evaluate the 12 potential analogs of F0317-0202 for broad-spectrum activity compared to their parent molecules, these analogs were screened against hookworms and whipworms using a dose-response of 0, 1, 3, 10, 30, and 90µM. Each dose was tested against 6 adult worms. Drug activity was determined using a worminator motility readout. GraphPad Prism generated the IC_50_ values utilizing the 48-hour average normalized motility of 6 individual worms.

## Authors contributions

M.A.E and R.V.A., conceived, designed, supervised, and interpreted the experiments. M.A.E and R.V.A. wrote the manuscript. M.A.E, Y.M.K., E.G., and P.C. performed the *in vitro* screening. M.A.E and S.S. performed data mining and analyzed SAR results. P.R.T. and L.B. contributed libraries, technical advice, and inputs on the manuscript. M.A.E. designed the graphic abstract. R.V.A designed the TOC graphic. R.V.A. secured funding.

## Supporting information

Supporting Information Available free of charge: Additional details for the potential nematode targets and additional experimental details for F0317-0202 tested analogs, chemical structures, and motility scores (Figure S1) (PDF).

## Notes

The authors declare the following competing financial interest(s): MAE and RVA are inventors on patents related to the compounds discussed in this manuscript, which have been filed by UMASS Chan Medical School. These patents cover the use of these compounds in the treatment of parasitic worm infections. No other conflicts of interest are reported.

## Acknowledgments

This work was financially supported by the National Institute of Health (USA) – National Institute of Allergy and Infectious Diseases grants R01 AI50866 and R01 AI056189 both to R.V.A.

## Notes

### Competing Interest Statement

The authors have declared no competing interest.

### Summary of Updates

New author, new experiments, and new references added. One new set of experiments have been added, which has added an author. References have also been updated.

