## Supporting Information A for "High-throughput screening of more than 30,000 compounds for anthelmintics against gastrointestinal nematode parasites"

**Additional details for potential nematode targets**

The 55 broad-spectrum compounds identified with potential activity against the adult stages of *A. ceylanicum* hookworm and *T. muris* whipworms were subjected to deep data mining, focusing on published activity against other eukaryotic organisms and, if available, especially for nematodes. Based on these analyses, we hypothesized potential nematode targets for some of these compounds based on their known mechanism of action in other organisms. Four examples are discussed below.

**BCL-2:**

Obatoclax was identified as a broad-spectrum hit. Obatoclax is a pan inhibitor of the pro-survival protein BCL-2. Upon inhibition of BCL-2, it leads to apoptosis and induction of BAX- and BAK-dependent and -independent cell death in many cancer cells, including hematological malignancies^1^. The BCL-2 proteins possess four BCL-2-homology domains (BH1, BH2, BH3, and BH4) that inhibit apoptosis via binding to the two sub-classes of pro-apoptotic proteins, BAX/BAK/BOK and BH3-only, which sense apoptotic stimuli and act as an activator in the process of BAX and BAK-mediated permeabilization of the outer mitochondrial membrane^1^. Obatoclax is classified as a BH3 mimetic that binds to the BH3 binding grove of the BCL-2, allowing the freed proapoptotic BH3-only protein to induce cellular death. Independent of inhibiting pro-survival BCL-2, obatoclax was also found to modulate BAX and BAK proteins to induce apoptosis. Obatoclax showed promising results in phases I and II and has reached phase III of clinical trials to treat hematological cancers and solid tumors^1^. BCL-2 protein homologs have been identified in 12 free-living nematodes and 60 animal parasitic nematodes in clades I, III, IV, and V of phylum Nematoda, with significant variation in BCL-2-like proteins within each taxon^2^, making BCL-2 a suitable candidate for designing parasite-selective inhibitors.

**PTPMT1:**

We identified alexidine as a broad-spectrum hit against *A. ceylanicum* hookworms and *T. muris* whipworms. Alexidine is a di-biguanide compound that is a known selective inhibitor of the Protein Tyrosine Phosphatase localized to Mitochondrion 1 (PTPMT1), which plays a critical role in cell signaling^3^. Treatment of rat pancreatic beta cells with alexidine affected the phosphorylation of mitochondrial proteins in a similar manner to the genetic knockdown of PTPMT1, and the knockdown of the same protein in rat islet cells makes them insensitive to the alexidine^3^. *In vivo* screening in zebrafish larvae identified alexidine as a glucose-lowering agent associated with an increased level of succinate dehydrogenase activity, a known PTPMT1 substrate; similarly, a mutation in *ptpmt1* eliminated the effect of alexidine in lowering glucose levels and resulted in hyperphosphorylation and activation of succinate dehydrogenase^4^. Structural and functional analysis of PTPMT1 demonstrated that loops essential for phosphatase function is highly conserved in human, mouse, chicken, zebrafish, insects, nematodes, and plants^5^, making it a suitable candidate for a pan-anthelmintic after addressing selectivity.

**NOX and NOS**

We identified diphenyleneiodonium as a broad-spectrum hit against *A. ceylanicum* hookworms and *T. muris* whipworms. Diphenyleneiodonium is a known inhibitor of NADPH oxidase (NOXes) was also found to strongly inhibit nitric oxide (NO) synthase (NOS)^6, 7^. NOS functional activity and NO-derived activity have been detected in the nervous structures of the *Ascaris suum* ^8^, *Toxocara canis* ^9^*,* and *Strongyloides venezuelenesis* ^10^. NOS is localized in the muscular wall of Brugia *malayi* and *Dirifilaria immitis* adult worms ^11, 12^. Helminth parasites, including nematodes, were found to induce NO production in their vertebrate hosts^10^. NO is an essential molecule involved in different biological functions of parasitic nematodes, most importantly neurotransmission, muscular relaxation, and detoxification of ROS^10^. ROS, whether generated from intrinsic (parasitic) or extrinsic sources (host), deleteriously impacts parasitic helminths^13^. The NOS enzyme may thus be an essential drug target.

**LXR:**

We identified T0901317 as a broad-spectrum hit against *A. ceylanicum* hookworms and *T. muris* whipworms. T0901317 is a well-known specific Liver X Receptor (LXR) agonist used to study the physiological effects of LXR^14-16^. LXRs (Nr1h2 and Nr1H3) are nuclear receptors that act as transcription factors regulating genes essential for cellular cholesterol uptake and lipid homeostasis^17^. LXRs are expressed in most tissues and are therapeutic targets in cardiovascular and neurological diseases^17^. LXRs are activated by various cholesterol derivatives and regulate the expression of genes associated with cholesterol homeostasis, such as the cholesterol 7α-hydroxylase gene (CYP7A) responsible for cholesterol removal and metabolism^18^. Nematodes can't synthesize cholesterol and rely on exogenous sources from the environment or their hosts^19, 20^. Thus, ligand activation of the LXRs ortholog in parasitic nematodes could result in cholesterol depletion and negatively impact parasite lipid homeostasis. In *C. elegans,* *daf-12* gene is the most like LXRs based on sequence similarity search, with functions linked to cholesterol and lipid metabolism^21^. Via gene expression analysis, *daf-12* regulated a network of genes involved in energy homeostasis and lipid metabolism; such a network was found to be conserved in the parasitic nematodes^22, 23^. T091317, tested at higher doses, showed deleterious effects on *C. elegans* *clk-1* mutant growth, possibly due to severe cholesterol depletion; however, lower doses decreased the defecation cycle length, and the same reduction in defecation cycle was observed in animals growing on cholesterol-reduced medium^15^.

**Additional details for F0317-0202 analogs chemical structure and anthelmintic activity**

| F0317-0202  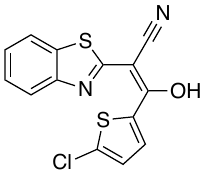  Hookworms 30µM Percent of Relative Motility (R.M.) : 1%  Whipworms 30µM Percent of Relative Motility (R.M.) : 0% | | | |
| --- | --- | --- | --- |
| F0317-0019  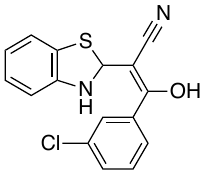  Hookworm R.M: 2%  Whipworms R.M: 22% | F0317-0013  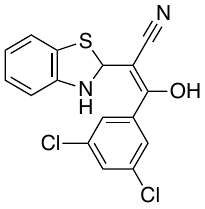  Hookworms R.M: 3%  Whipworms R.M: 0% | F0317-0161  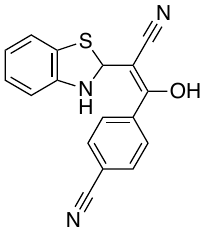  Hookworms R.M: 1%  Whipworms R.M: 2% | F0336-0103  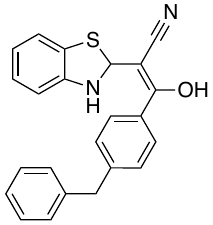  Hookworms R.M: 1%  Whipworms R.M: 50% |
| F0317-0175  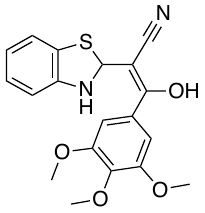  Hookworm R.M: 1%  Whipworms R.M: 37% | F0328-0210  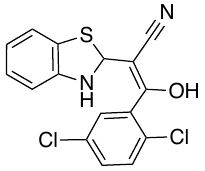  Hookworm R.M: 1%  Whipworms R.M: 8% | F0317-0012  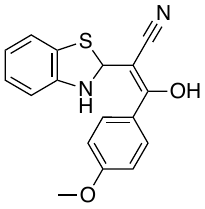  Hookworm R.M: 2%  Whipworms R.M: 16% | F0317-0009  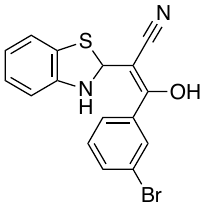  Hookworm R.M: 2%  Whipworms R.M: 1% |
| F0328-0550  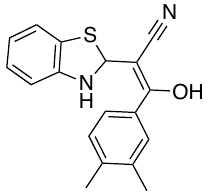  Hookworm R.M: 1%  Whipworms R.M: 20% | F0317-0160  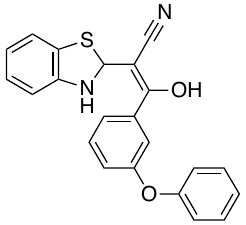  Hookworm R.M: 6%  Whipworms R.M: 44% | F0882-0289  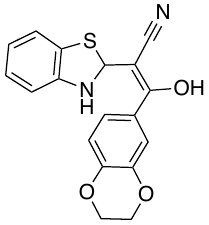  Hookworm R.M: 11%  Whipworms R.M: 42% | F0463-0179  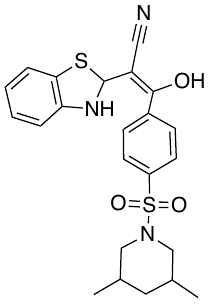  Hookworm R.M: 33%  Whipworms R.M: 1% |
| F0317-0186  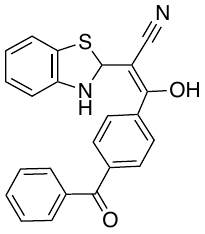  Whipworms R.M: 92% | F0466-0338  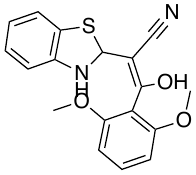  Whipworms R.M: 100% | F0317-0189  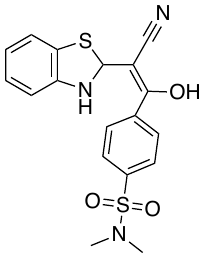  Whipworms R.M: 71% | F0463-0263  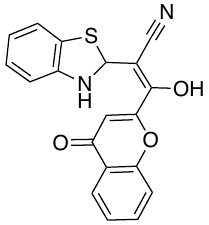  Whipworms R.M: 100% |
| F9995-0805  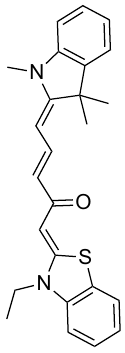  Hookworms R.M: 77%  Whipworms R.M: 62% | F9995-0794  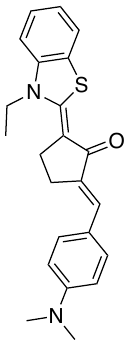  Hookworms R.M: 80%  Whipworms R.M: 79% | F1284-0734  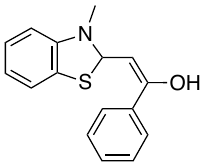  Whipworms R.M: 80% | F0363-0011  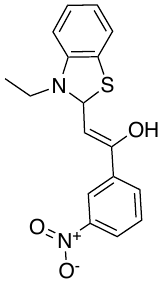  Whipworms R.M: 89% |
| F3351-0269  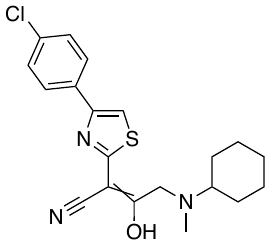  Whipworms R.M: 100% | F3055-0633  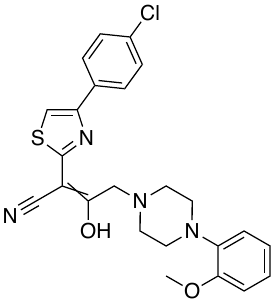  Whipworms R.M: 100% | F3375-2458  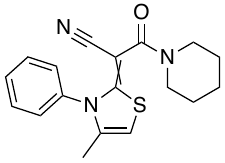  Whipworms R.M: 100% | F1589-1467  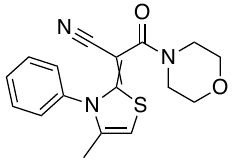  Whipworms R.M: 100% |
| F1589-1468  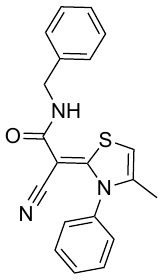  Whipworms R.M: 100% | F3077-0352  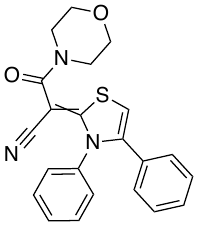  Whipworms R.M: 100% | F3077-0348  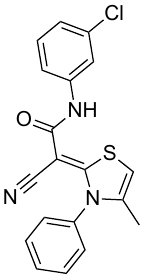  Whipworms R.M: 100% | F3055-0633  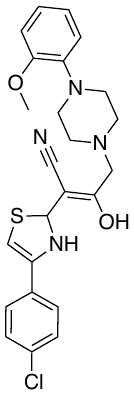  Whipworms R.M: 100% |
| **Figure S1:** F0317-0202 analogs structures and their activity against *A. ceylanicum* Hookworms and *T. muris* whipworms as percent of Relative Motility (R.M) to the DMSO control, recorded after 48 hours of exposure to 30µM (100% is healthy; 0% is non-motile). Two of the compounds (F9995-0805, F9995-0794) with poor activity against whipworms were also tested against hookworms to confirm that low whipworm activity compounds were not highly active against hookworms. | | | |
